# Elucidating the effect of cations on *Klebsiella pneumoniae* biofilm and related genes

**DOI:** 10.1101/2024.01.24.577145

**Authors:** Aravind Murugavel, Srinithi Gunasekaran, Jayapradha Ramakrishnan

**Affiliations:** Actinomycetes Bioprospecting Lab, Centre for Research in Infectious Diseases (CRID), School of Chemical and Biotechnology, SASTRA University, Tirumalaisamudram, Thanjavur -613401, Tamilnadu, India; Actinomycetes Bioprospecting Lab Centre for Research in Infectious Diseases (CRID), School of Chemical and Biotechnology, SASTRA University, Tirumalaisamudram, Thanjavur -613401, Tamilnadu, India

**Keywords:** *Klebsiella pneumoniae*, Urinary tract infection, Biofilm, capsule, fimbriae, Lipopolysaccharide

## Abstract

*K. pneumoniae* is a non-motile, encapsulated bacterium from the Enterobacteriaceae family. The illnesses related to this opportunistic pathogen are pneumonia, Urinary Tract Infection (UTI), pyogenic liver abscess, endophthalmitis, and meningitis. Among them, UTI is predominant due to its biofilm formation leading to the mortality of 150 million people worldwide. The function of monovalent and divalent ions on *Klebsiella* biofilm, aside from physiochemical variables, remains unclear. Hence the present study was performed to analyze the role of K^+^, Ca^2+^, Na^+^, Mg^2+^, NH^+^_4_ in biofilm formation and its influence on biofilm-related genes. Among the tested cations, K^+^ and Ca^2+^ yielded strong biofilm in clinical and environment isolates in pH between 6.5 to 9.5. Increasing Ca^2+^ ions concertation reduced the *Klebsiella* biofilm. When compared to the hypercalciuria condition (Ca^2+^ level > 5 mM), 2.5 mM resulted in high biofilm mass. Cations concurrently enhanced the size of the capsule and cell density of *Klebsiella* but were not correlated with biofilm mass. Expression of the LPS gene (*WbaG*) either in planktonic or biofilm stage promoted biofilm formation in the presence of K^+^, Ca^2+^ and Na^+^. Whereas, expression of fimbriae genes (*FimH* and *mrkD*) was co-regulation, and capsule genes (*RmpA* and *Wcab*) were absent. Stating, the primary component needed for the *Klebsiella* biofilm is not the capsule or fimbriae, rather LPS. Resulting as a potent target in the treatment of *Klebsiella* biofilm in UTI. To the best of our knowledge, this is the first kind of study on the effect of cation ions on biofilm and planktonic cells of *Klebsiella* spp. and demonstrating the role of LPS biosynthesis gene (*WbaG*) in biofilm development.

## Introduction

*Klebsiella pneumoniae* is a potential opportunistic pathogen with the significant virulence factors such as capsular polysaccharide, LPS, type 1 and type 3 fimbriae and siderophores. The amendment of this virulence factors regulates the strain more virulent. Such as antiphagocytic polysaccharide capsule and fimbriated strain inducing high hydrophobic mediating adherence to the abiotic surface & host tissue (Ruiz et al. 1993, X. Huang et al. 2022, Di Martino et al. 2003b). In our recent investigation, comparing to planktonic cells, the biofilm interferes with innate immunity and reduces phagocytic activity (Sudarshan Singh Rathore et al. 2022). This emphasising the importance of biofilm development with its major components. Such as capsule, LPS and fimbriae as a significant feature for its virulence and establishment of persistent infections. Causing central venous catheter-related blood stream infections, both severe and uncomplicated urinary tract infections, endotracheal tube colonisation, and ventilator-associated pneumonia. These infections are significantly associated with morbidity and mortality world-wide. Among the various virulence factors, biofilm formation is a critical factor in the development of bacteraemia. Biofilms assist pathogens in evading phagocytosis and are the most common cause of bloodstream infections. They are resistant to antimicrobial therapies, resulting in therapeutic failure causing significant threat, globally. This underlines the importance of a better knowledge on *K. pneumoniae* biofilm production and pathogenesis. The elements that contribute to *K. pneumoniae* biofilm, role of polysaccharide capsule, LPS, fimbriae and pili, and iron metabolism, have been thoroughly investigated to develop efficient treatment techniques. Apart from the cellular components, environmental factors also play an important role in biofilm formation. Various other physiochemical parameters such as substratum properties, effect of temperature, pH, oxygen level, urea and ion concentration were evaluated. The role of divalent and monovalent ions on *Klebsiella* biofilm which may impact the biofilm properties were not elucidated.

Hence to control biofilm-related illnesses, a fundamental understanding of *K. pneumoniae* biofilm development is required. Furthermore, cation ions and cation homeostasis have been extensively researched in planktonic microorganisms. Cation ions such as Zn (II), Mn (II) and Fe between 0.4-1 mM favoured bacterial growth. On the other hand, starvation, deficiency and excess of cation ions inhibit the bacterial growth (McDevitt et al. 2011, Gonciarz and Renslo 2021). Such kind of extensive studies on effect of cation ions on *Klebsiella* biofilm development is lacking. As biofilm cells are quite distinct from planktonic cells in terms of gene expression and survival ability. The effect of cation ions on biofilm formation in some bacterial species, such as *B. subtilis, S. aureus, P. fluorescens and Exigupbacterium* sp. has been investigated (Sharipova et al. 2023), Vaezi et al. 2021, Song and Leff et al. 2006, Pavez et al. 2023). However, its influence on *Klebsiella* sp. biofilm formation and related genes has not been investigated. Such a correlation with vital biofilm genes could result in important discoveries. The influence of cation ions on biofilm growth was investigated in this work using several strains of *Klebsiella* sp. of clinical and environmental origin. In addition, to verify the responsiveness of biofilm-related genes such *WcaG* (Capsule biosynthesis), *RmpA* (Regulator of mucoid phenotype, *WabG* (LPS biosynthesis), and *FimH* & *mrkD* (Adhesin) was also investigated. To the best of our knowledge, influence of divalent and monovalent ions on biofilm formation and expression of biofilm related genes of *Klebsiella* sp. is of first kind.

## Methodology

### Strains used in this study

A total of seven *K. pneumoniae* strains were tested, namely four uropathogenic (kp1,kp2,kp3) isolates collected from the D.Y. Patil hospital and research centre in Pune, two blood isolates (BA 33569 and BA 28434) from Christian Medical College (CMC) Vellore, Environmental (kp4) and clinical reference strain (kp5) from MTCC-Chandigarh. For presumptive identification, all strains were grown on the Hi-Crome UTI agar medium (Lalitha et al. 2017)

### Biofilm development with varying Cation ion at 72 h

To study the effect of various cation ions, the strains were cultured in LB broth at 37 ^0^C, 24 h. Centrifuged at 7000 g for 10 minutes and cell pellet were washed with sterile distilled water thrice, resuspended in same. The working stock culture was adjusted to 0.08 OD_595_ equivalent to ∼10^8^ CFU ml^-1^. In a sterile 96 well polystyrene microtiter plate, 10 µl volume of working stock culture was added to 190 µl of various ions and control glucose (2 mM) medium. The concentration of cation ions in the form of salt was employed in relation to Artificial Urine Medium (AUM) reported by (Brooks and Keevil 1997) (Suppl.Table 4). The ion medium contains 2 mM glucose along with the salts such as (MgSO_4_.7H_2_O-2 mM), (NaHCO_3_ -25 mM), (KH_2_PO_4_ -7 mM), (NH_4_Cl-25 mM) and (CaCl_2_.H_2_O-2.5 mM). After 72 h of incubation at 37 ^0^C, the wells were washed thrice with PBS and dehydrated in an inverted position. Using 50 µl of 0.1% crystal violet, the biofilm mass was stained and washed thrice in 200 µl of distilled water. The stained biofilm mass was dissolved in 200 µl of 33% glacial acetic acid. At 595 nm, optical density (OD) was measured using a microplate reader.

Similarly, influence of pH and concertation of Ca^2+^ ion on biofilm was carried. Using HCL and NaOH, the pH was adjusted from 4.5 to 9.5. The concertation of Ca^2+^ ion was increased in two-fold interval to maximum of 7.5 mM.

### To observe the correlation of colony forming unit, capsule size measurement and colony morphology characteristics with biofilm

To observe the capsule size, colony forming unit and morphology characteristics, all the strains were adjusted with 0.8 OD ∼ 1x10^8^ CFU ml^-1^ in all respective salt medium carried in biofilm assay. For CFU analysis,10 µl of the sample was serially diluted to 10^5^ and the final volume of 10 µl layered on respective salt agar medium (Mg^2+^, K^+^, NH^+^_4_, Na^+^, Ca^2+^, AS). After 24 h, CFU was calculated.

For capsule size determination, 20 µl volume of sample is equally mixed with nigrosine strain and observed under Nikon eclipse ci light microscope in 1000x magnification and images were exported using Nikon digital camera D5100. To differentiate the capsule size (Dwc) and cell body weight limited by cell wall (Dcb), the images was optimized using ImageJ 1.48v software (National Institutes of Health, Washington, DC, USA) following guidelines of Best Practice Guidelines on Publication Ethics (Graf et al. 2007). The size of the capsule relative to that of the whole cell was defined, as a percentage as {[(Dwc – Dcb)/Dwc] × 100}. Ten cells were measured for each determination and average was calculated (S. Rathore et al. 2016).

Colony morphology, 10 µl volume of the sample was spotted petri plate containing respective salt ion with addition of 0.8 mg L^-1^ of congo red and observed after 72 h.

### Fluorescence imaging

To further confirm the biofilm quantification of CV assay, fluorescence imaging was carried out. In a sterilized glass slide (22 mm x 22 mm) placed in 6 well polypropylene plates. 150 µl of culture containing 1×10^8^ CFU ml^-1^ were inoculated into 3 ml of respective salt medium. After 72 h incubation, the slide was washed thrice using PBS and left for air dry. The slides were strained with 20 µg ml^-1^ of fluorescein isothiocyanate con A (FITC con-A) and observe under fluorescence microscope Nikon eclipse ts2.

### mRNA isolation from planktonic cells and biofilm mass

Based on the biofilm formation ability and difference in source, Strong Biofilm Producing (SBP) strain “kp3” and Lowest Biofilm Producer (LBP) “BA 28434” were carried for gene expression in response to all conditions. The mRNA isolation protocol was referred (Atshan et al. 2012) technique was consulted for the mRNA isolation procedure for both planktonic and biofilm cells.

### cDNA conversion and Real time PCR

The quality and purity of isolated mRNA were assessed by NanoDrop and Gel electrophoreses. Using PrimeScript™ RT Reagent Kit (Perfect Real Time) (Cat no: RR037A) with given the standard protocol the cDNA was performed. For the gene expression analysis, Quantstudio 5 and TB Green® Premix Ex TaqTM II (Tli RNase H Plus) (Cat no: RR820A) were employed. To the total of 10 µl of reaction mixture, 5 µl of TB Green Premix Ex Taq II (Tli RNaseH Plus) (2X), 1µl of (0.5mM) of both forward and reverse primers (Table 1), 1µl (100 ng) of template cDNA were added. Cycling program include of initial denaturation at 95 ^0^C for 30 s for enzyme activation, denaturation at 95 ^0^C for 5 s, annealing temperature (Table 1) for 45 s and extension for 1 min at 72 ^0^C, followed for default melting temperature and carried for 45 cycles. Using Relative Expression Software Tool (REST), the comparative gene expression was measured (Pfaffl, Horgan, and Dempfle 2002)

**Table 1:**
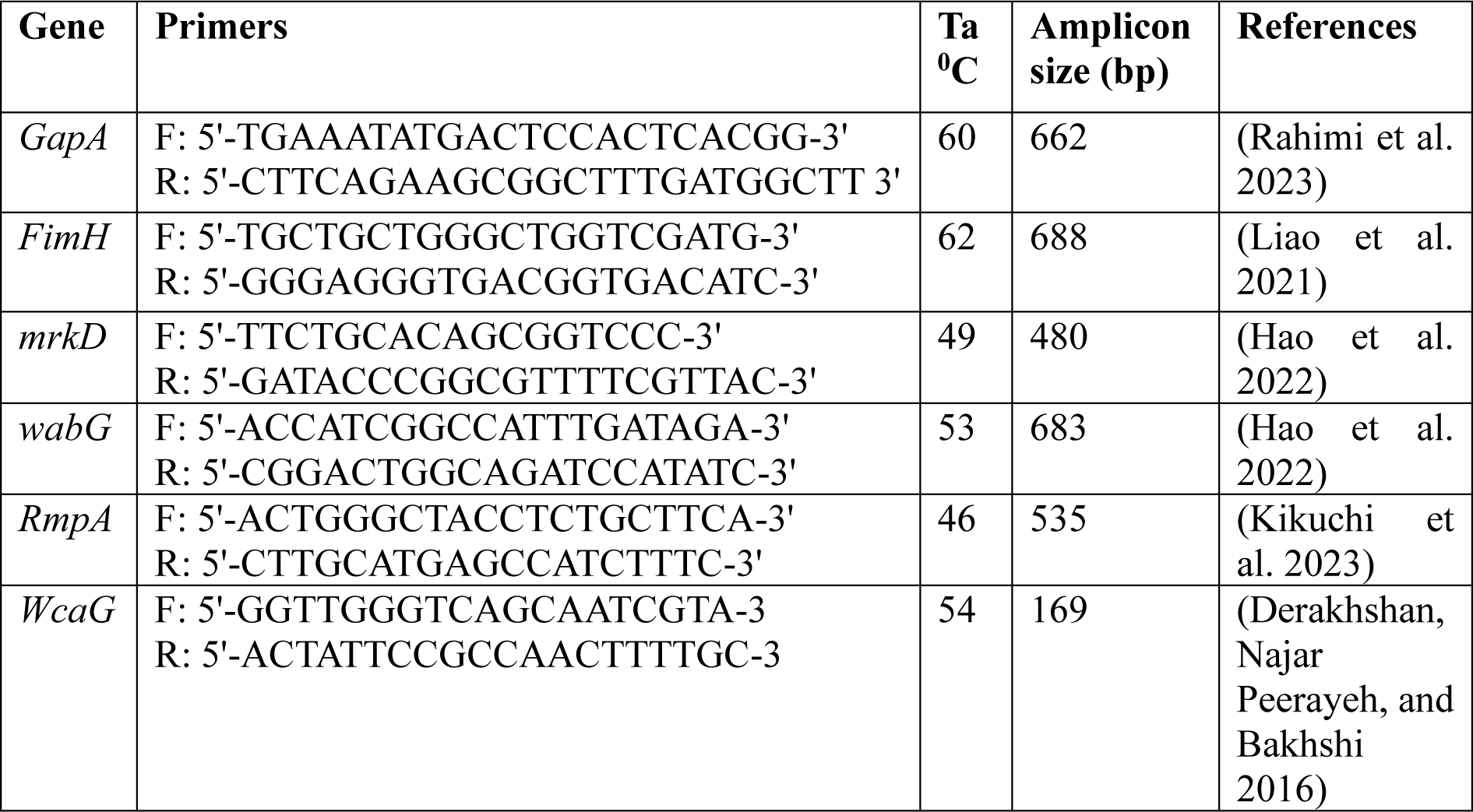
Primers with its annealing temperature and amplicon size (bp)

### Statistical analysis

All the tests were carried out in two independent trails with three replicates. The data represented in the graph and analysis were performed using GraphPad Prism version 8. The significance difference was calculated using two-way ANOVA with Bonferroni post-test. The data represent mean Standard Deviation (SD) is calculated for significance differences value of p<0.001.

## Results

### Ca^2+^ and K^+^ ions promote the development of biofilms in *Klebsiella* sp

We analysed the effect of various cation ions for biofilm growth of seven different *K. pneumoniae* strains of clinical and an environmental origin. When comparing the biofilm growth in glucose (2 mM) supplemented with different cation ions, the highest order of biofilm formation for the tested strains are as follows: BA 33560 - 66% (K^+^); kp3 strain - 65% (Ca^2+^); kp2 - 57% (Na^+^,); kp1- 56% (Ca^2+^); kp5 - 50% (AS); BA 28434 - 48% (Ca^2+^); kp4 -43% (K^+^) (Figure 1).

**Figure 1:**
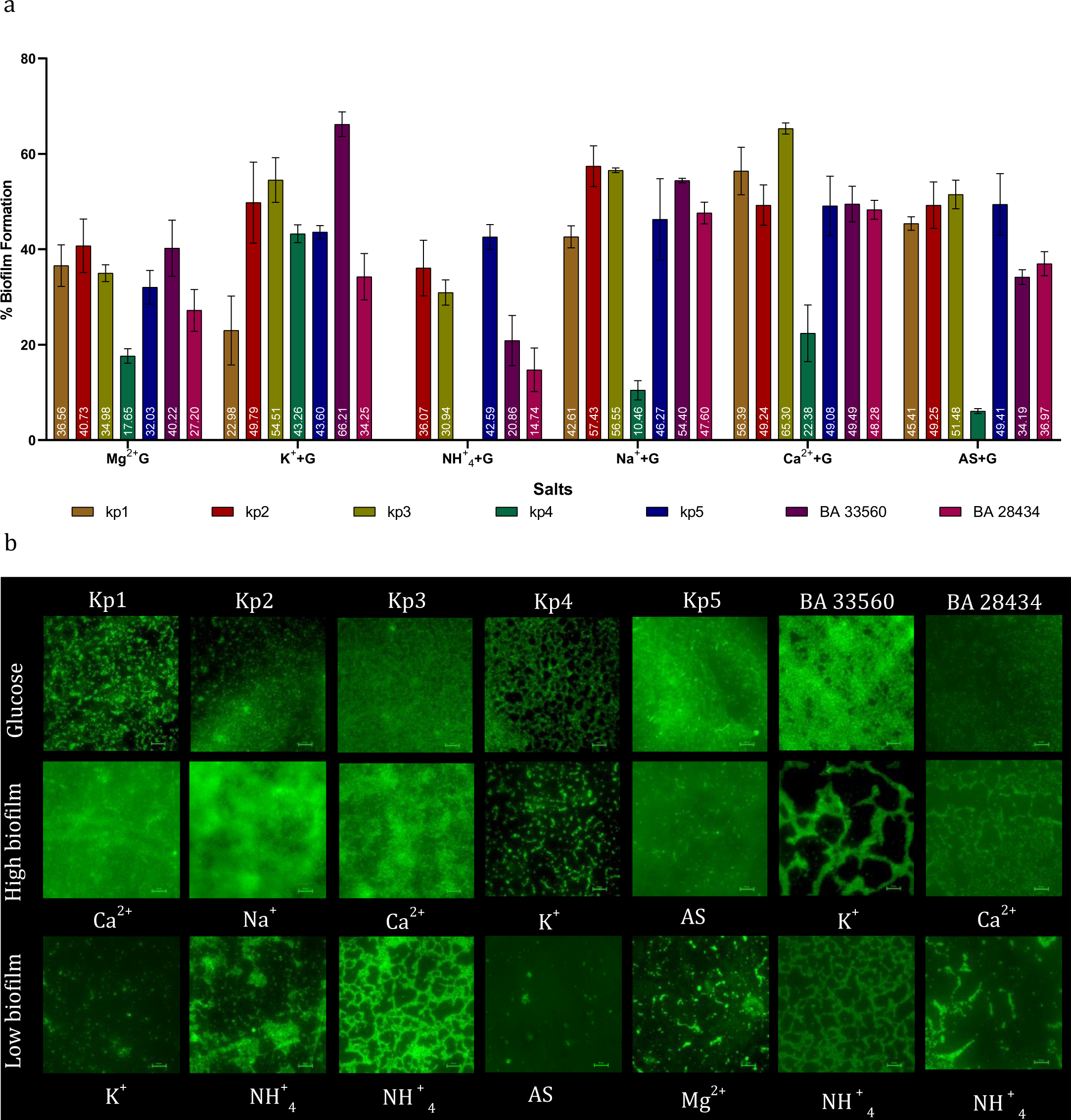
A) *K. pneumoniae* biofilm formation in the presence of Glucose, Mg^2+^, K^+^, NH^+^_4_, Na^+^, Ca^2+^ and AS incubated for 72 h at 37 ^0^C. Ca^2+^ and K^+^ followed with Na^+^ showed the highest biofilm development in seven distinct strains, including environmental and clinical strains. The bar displays the mean ± SD for (n = 3) with *p <0.001. B) Fluorescence imaging of two biofilm conditions—high and low—for seven distinct strains of *K. pneumoniae* (urine isolates (kp1, kp2, kp3, kp4, kp5), Blood isolates (BA 33560 & BA 28434) and environment isolate (kp4).

Ca^2+^ and K^+^ ions enhanced biofilm density followed by Na^+^ in all isolates (urine, blood and environmental). Stating Ca^2+^ and K^+^ ions significantly increase the development of biofilm in *Klebsiella* sp. regardless of source (*p* < 0.001). The most prevalent cation required by every living cell is K^+^, similarly the present research highlights its significant role in biofilm formation of *K. pneumoniae*. The divalent cations, such as Ca^2+^ and Mg^2+^ influence biofilm through electrostatic interactions. However, in some species, their effects are dose dependent and debatable. For instance, *S. aureus* and *B. subtilis* respectively displayed lower biofilm thickness and topography at higher Ca^2+^ and Mg^2+^ concentrations (Sudhir K. Shukla, T. Subba Rao 2013). Indicating the effects of Ca^2+^ and Mg^2+^ ion on biofilm growth is independent between species. However, we found Ca^2+^ ion enhanced biofilm of *Klebsiella* sp. Similarly, in mucoid *P. aeroginosa*, Ca^2+^ ion regulated ten times thicker biofilm (Sarkisova et al. 2005). In case of *Bacillus* sp. presence or absence of Ca^2+^ did not impede the biofilm matrix reported by Masaki Nishikawa and Kazuo Kobayash et al., but significantly weakened biofilm architecture (Nishikawa and Kobayashi et al. 2021.). Similarly, the plant pathogen *Xylella fastidiosa* had no discernible impact on the levels of Exopolysaccharides (EPS) (Cruz et al. 2013). Our findings imply that Ca^2+^ and Mg^2+^ ions have a favourable impact on the development of *Klebsiella* sp. biofilm. Like *S. aureus* and *Closteridium* sp., Na^+^ ions also aided in the development of *Klebsiella* biofilm in our study (Vaezi et al. 2021, Philips et al. 2017). The biofilm is hindered or diminished when NH^+^4 ions are present. The medium containing combination of all ions has no discernible impact on the development of *Klebsiella* biofilms. In conclusion, *K. pneumoniae* biofilm growth is significantly impacted by Ca^2+^ and K^+^.

### *Klebsiella* biofilm development is stable in an alkaline environment

We investigated the effect of pH for the biofilm growth of *Klebsiella* sp. To study the effect of pH, the Glucose + Ca^2+^ media with different pH from 4.5 to 9.5 with an interval of 1 and pH 6.5 as control. According to the data, biofilm production was similar for pH values between 6.5 and 9.5 and significantly reduced for pH values below 4.5. (Figure 2). This is highly concurrent with the tolerance range of planktonic cells of *Klebsiella* sp. (pH 8 to 10).

**Figure 2:**
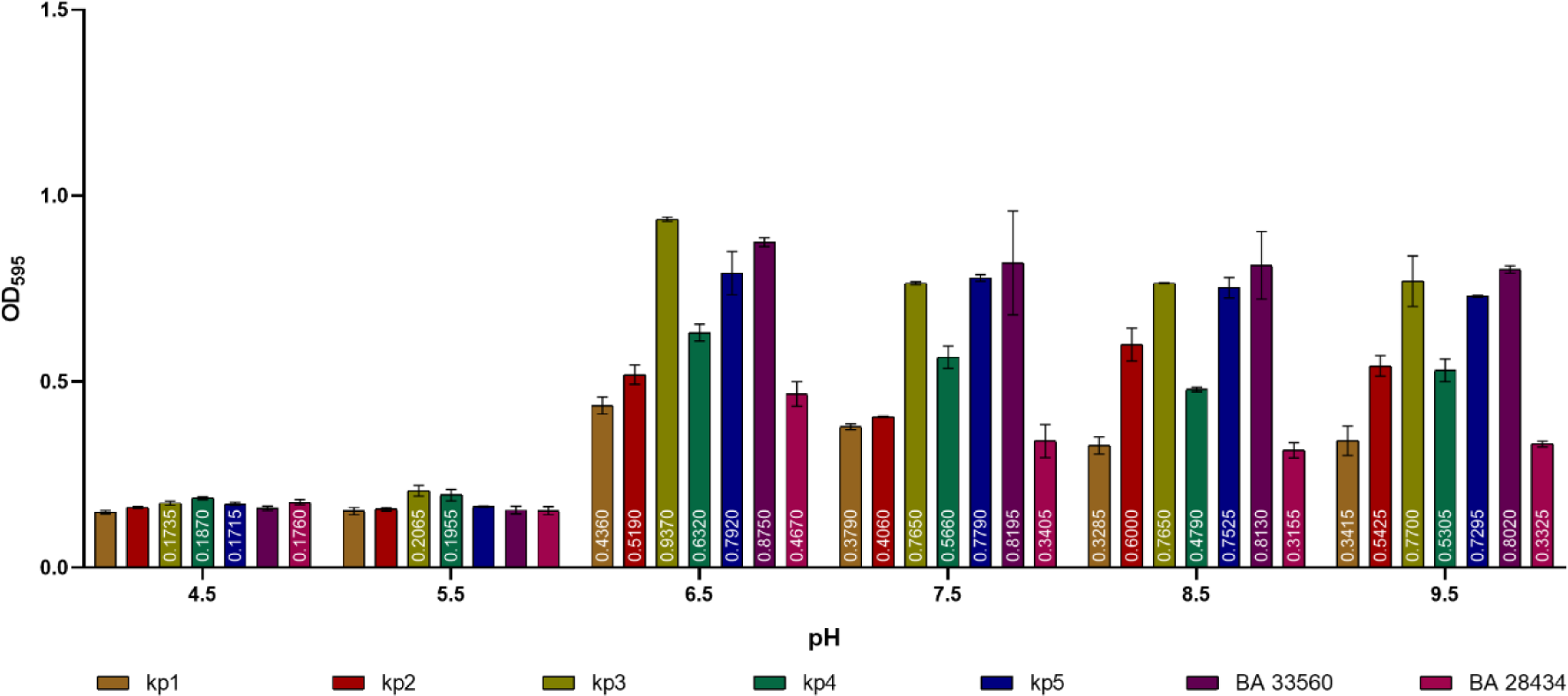
Biofilm development of *K. pneumoniae* in Ca^2+^ ions at varying pH settings. An acid pH (<6.5) displayed less biofilm development. Alkaline and neutral pH ranges of 6.5 to 9.5 moderated the growth of biofilms in a negligible way. The bar displays the mean ± SD for n = 3, with *p <0.001.

### Increase in Ca^2+^ concentrations decrease biofilm formation of *Klebsiella* sp

Ca^2+^ and K^+^ ions both significantly accelerated the formation of biofilms. Furthermore, we chose Ca^2+^ ions to continue with to better understand the effects of different concentrations. Ca^2+^ was tested from 3.5 mM to 7.5 mM with an interval of 1 mM. When compared to control (2.5 mM Ca^2+^+ 2 mM Glucose), all the strains of *Klebsiella* sp. displayed decrease in biofilm biomass except for kp1 and BA 28434 (Figure 3). In these two strains biofilm production was enhanced notably at 3.5 mM and decreased afterward (Figure 3). Similarly, the biofilm formation of *Citrobacter werkmanii* BF-6 (400 mM) and *Cronobacter sakazakii* (22 mM) were enhanced notably, but the enhancement presented a non-monotonic trend (Zhou et al. 2016, Ye et al. 2015). In case of *B. subtilis*, biofilm stabilized between 0.4 to 1.7 mM the concentration of calcium ion (Nishikawa and Kobayashi et al. 2021). The oral biofilm increased double the time when the medium was supplemented with 5 mM CaCl_2_ and reported that Ca^2+^ in the range of 16–64 mM, the biofilm biomass of *Pseudomonas* and *Arthrobacter* sp. were increased sharply in a concentration-dependent manner (Shokeen et al. 2022). In *P. fluorescens*, the biofilm biomass was increased at Ca^2+^ concentration up to 1 mMol L^-1^, and decreased greater than 10 mMol L^-1^ (Jiemin et al. 2019). These results suggested that the effect of various concentrations of Ca^2+^ ions varied with bacterial species. Hence a strain specific dose dependent study is required to understand the role of Ca^2+^ ions in *Klebsiella* sp. The study includes 2.5 mM Ca^2+^ ions which reflected a normal urine concentration, while higher than 5.5 mM Ca^2+^ concentration was reported as hypercalciuria. The concentration study demonstrates that normal urine conditions favour biofilm formation of *Klebsiella* sp. rather than hypercalciuria.

**Figure 3:**
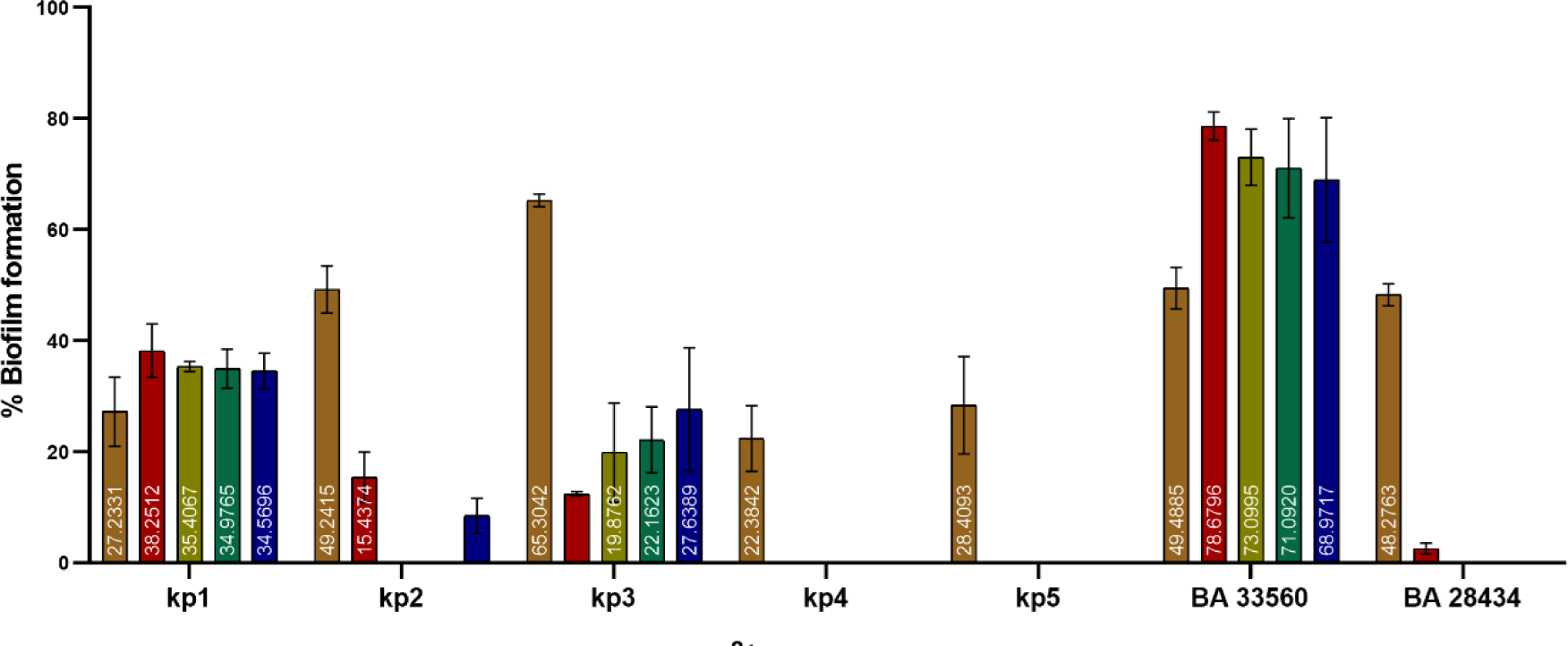
Biofilm development of *K. pneumoniae* at different Ca^2+^ ion concentrations, incubated for 72 h at 37 ^0^C. Except for BA 33560, all six strains showed that 2.5 mM was the ideal concentration for high biofilm development. The bar displays the mean ± SD for n = 3, with *p <0.001.

### Biofilm production is unrelated to cell counts

To confirm the effect of ions specifically on biofilm development, planktonic cells were estimated and compared with biofilm quantification . The strains were grown in the selective ion medium for 24 h and plated on cation medium supplemented with agar. When compared to control (Glucose 2 mM), the cell counts were found to have an average of 3.8-fold increase in the medium supplemented with ions. The increment in cell numbers is as for all the strains are as follows: All salt (5.2-fold), K^+^ (4.3-fold), NH^+^_4_ (3.8-fold), Ca^2+^ (3.6-fold), Na^+^ (3-fold), and Mg^2+^ (2.8- fold) increased cell populations (Table 2). The cell counts of the blood isolates were nearly identical to control and had no influential effect on planktonic cells. We observed that the relationship between ions impacts on the quantification of planktonic cells and biofilms is inverse. For instance, kp3 with 8.5-fold cell populated exhibited 65% biofilm when compare to 51% of AS biofilm with 12.8-fold cell density. The similar findings were displayed in other strains. The inverse proportion unequivocally demonstrates the crucial role Ca^2+^, K^+^, and Na^+^ ions in the formation of *Klebsiella* biofilm.

**Table 2:** Colony forming unit (CFU) of seven strains.

| Strain | Cation ion | CFU fold change<br>with respect to<br>Glucose | Biofilm<br>Percentage |
| --- | --- | --- | --- |
| **Kp1** | Mg<sup>2+</sup> | 5.4615 | 36% |
|  | K<sup>+</sup> | 6.9231 | 22% |
|  | NH<sup>+</sup><sub>4</sub> | 4.1692 | 0% |
|  | Na<sup>+</sup> | 6.1538 | 42% |
|  | Ca<sup>2+</sup> | 3.8923 | 56% |
|  | AS | 15.385 | 45% |
| **Kp2** | Mg<sup>2+</sup> | 3.3333 | 40% |
|  | K<sup>+</sup> | 8.333333333 | 49% |
|  | NH<sup>+</sup><sub>4</sub> | 6.6667 | 36% |
|  | Na<sup>+</sup> | 3.8889 | 57% |
|  | Ca<sup>2+</sup> | 6.6667 | 49% |
|  | AS | 4.4444 | 49% |
| **Kp3** | Mg<sup>2+</sup> | 7.6923 | 34% |
|  | K<sup>+</sup> | 7.9915 | 54% |
|  | NH<sup>+</sup><sub>4</sub> | 7.6923 | 30% |
|  | Na<sup>+</sup> | 6.8376 | 56% |
|  | Ca<sup>2+</sup> | 8.547 | 65% |
|  | AS | 12.821 | 51% |
| **Kp4** | Mg<sup>2+</sup> | 13.426 | 17% |
|  | K<sup>+</sup> | 15.404 | 43% |
|  | NH<sup>+</sup><sub>4</sub> | 21.277 | 0 |
|  | Na<sup>+</sup> | 12.766 | 10% |
|  | Ca<sup>2+</sup> | 17.021 | 22% |
|  | AS | 12.766 | 6% |
| **Kp5** | Mg<sup>2+</sup> | 1.0769 | 32% |
|  | K<sup>+</sup> | 3.4874 | 43% |
|  | NH<sup>+</sup><sub>4</sub> | 2.0588 | 42% |
|  | Na<sup>+</sup> | 3.3613 | 46% |
|  | Ca<sup>2+</sup> | 2.7311 | 49% |
|  | AS | 4.2017 | 49% |
| **BA 33560** | Mg<sup>2+</sup> | 1.3011 | 40% |
|  | K<sup>+</sup> | 3.719 | 66% |
|  | NH<sup>+</sup><sub>4</sub> | 4.1322 | 20% |
|  | Na<sup>+</sup> | 1.0083 | 54% |
|  | Ca<sup>2+</sup> | 2.4793 | 49% |
|  | AS | 2.4793 | 34% |
| **BA 28434** | Mg<sup>2+</sup> | 1.0596 | 27% |
|  | K<sup>+</sup> | 2.2098 | 34% |
|  | NH<sup>+</sup><sub>4</sub> | 1.51 | 14% |
|  | Na<sup>+</sup> | 4.1182 | 47% |
|  | Ca<sup>2+</sup> | 2.0223 | 48% |
|  | AS | 1.3245 | 36% |

### Colony morphology is unrelated to *Klebsiella* biofilm

To further confirm the role of ions on biofilm, we investigated the effect of ions on colony morphology and its relation with biofilm biomass. The description of colony morphology on each ion was provided in (Suppl.Table 3), which demonstrates that each ion had a unique morphotype common to all 7-strains (Figure 4b) and was not co-related to *Klebsiella* biofilm development. In contrast to the divalent cation ions (Ca^2+^ & Mg^2+^), monovalent cation ions (K^+^, NH^+^_4_ & Na^+^) induced pigments in all tested strains. Whereas, in presence of K^+^ ions, BA 28434 and kp1 produced black colour. In Na^+^ ion medium, BA33560, BA 28434, kp1 and kp3 appeared as pink colour. In NH^+^_4_ ion condition, kp5 & kp3 strain also produced pink colour. Whereas kp2, BA 28434, kp1, kp4 strains appeared red colour. Other than pigment, no other discernible similarity was found.

**Figure 4:**
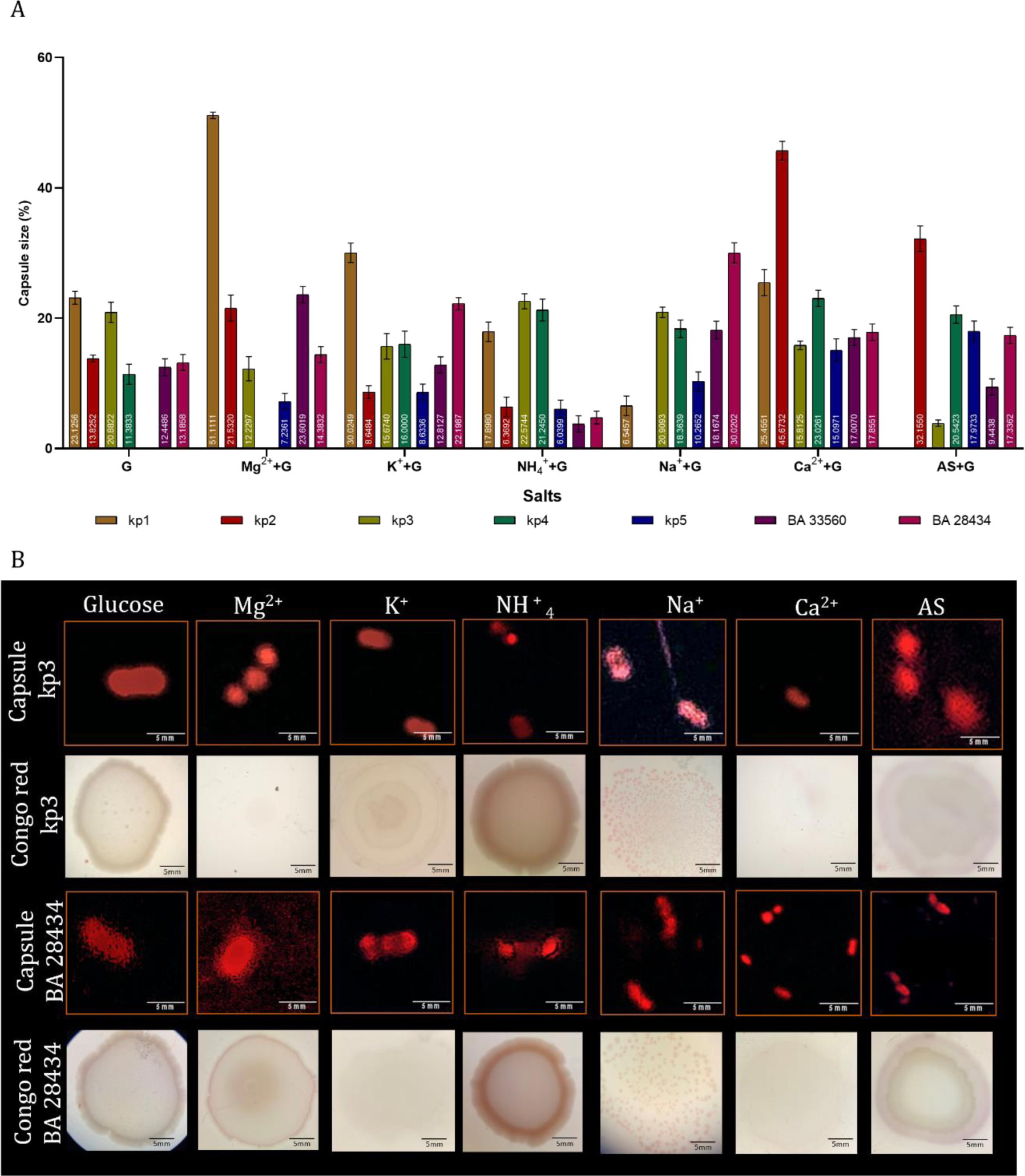
A) *K. pneumoniae* capsule formation in the presence of Glucose, Mg^2+^, K^+^, NH^+^_4_, Na^+^, Ca^2+^ and AS. All strains had independent capsule sizes regardless of the presence of cations. The bar represent mean ± SD for (n = 8) with *p<0.001. B) Capsule imaging and Biofilm morphology of *K. pneumoniae* (kp3 and BA 28434) strains.

### Capsule thickness has less impact in biofilm formation

By measuring the capsule size of planktonic cells cultured in various ion mediums, the role of ions in capsule induction and its influence on biofilm development was studied. We were able to investigate how different ions influence capsule growth by comparing the cell size and capsule thickness. The highest biofilm producer of kp3 strain in presence of Ca^2+^ induced 65% of biofilm production with capsule size of 15%. Parallelly, in presence of All salt (AS), with 3% capsule size resulted 51% of biofilm formation. A similar observation was observed for other strains as well (Figure 4(a-b)). These findings imply that there is no relationship between capsule thickness and biofilm biomass. Believing that increasing capsule size does not favour biofilm production suggesting the less role of encapsulation in biofilm formation of *Klebsiella* sp.

### qPCR Gene expression

#### rmpA gene

The kp3 (SBP) and BA 28434 (LBP) strains were selected among the tested strains to examine the expression of biofilm-related genes in planktonic and biofilm cells under various salt conditions. Apart from all salts of SBP strain, *rmpA* expression was downregulated in the planktonic cells of both strains. In the presence of Mg^2+^, it was discovered that both strains had overexpression in biofilm cells of 8.5-fold and 6-fold, respectively. However, the agarose gel result was not found to have the *rmpA* amplicon (488 bp) (Figure 5) . These findings imply that there was no *rmpA* expression in both the strains in planktonic and biofilm.

**Figure 5:**
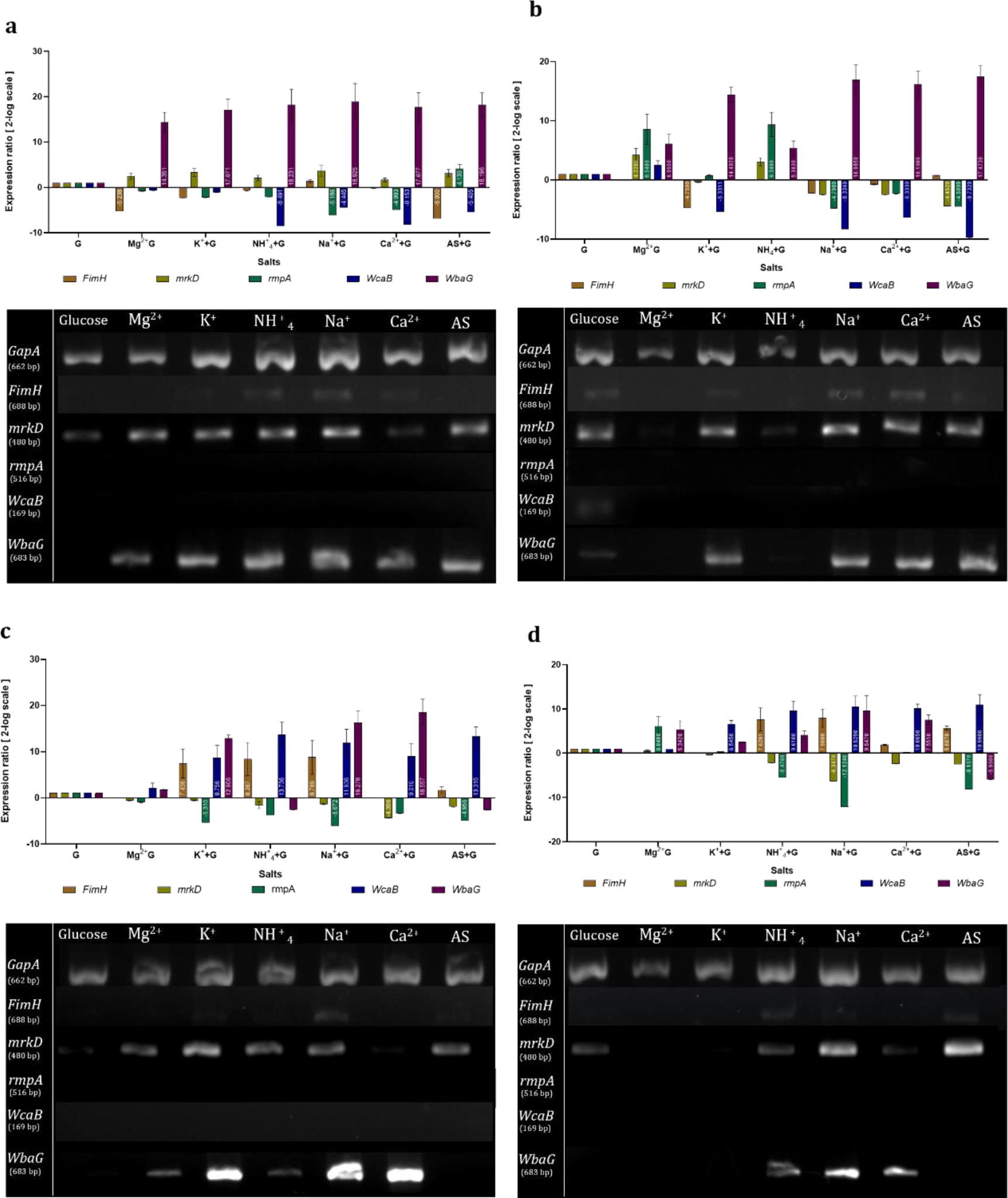
Plot shows the relative expression ratio and gel electrophoresis of real-time gene expression in the presence of Glucose, Mg^2+^, K^+^, NH^+^_4_, Na^+^, Ca^2+^, and AS for the genes *GapA, FimH, mrkD, rmpA, Wcab,* and *WbaG*. a) kp3 strain- Planktonic cells; b) kp3 strain-Biofilm cells; c) BA 28434-Planktonic cells; d) BA 28434-Biofilm cells of *Klebsiella pneumoniae* strains. The bar displays the mean ± SD for (n = 2) with *p <0.001.

We also noticed that *rmpA* expression and capsule size are not related. For example, in the presence of Mg^2+^ and K^+^, the capsule size of SBP strain increased by 12% and 15%, respectively, but *rmpA* expression downregulated. A similar correlation was observed in BA 28434 strain. Several studies have found that *rmpA/rmpA 2* positively regulate Capsular Polysaccharide (CPS) gene expression in *K. pneumoniae*. In our investigation, we found no such association in terms of capsule size. One possible explanation could be the expression of *rmpA2*, which we did not investigate.

#### FimH gene

*FimH* was down regulated in the planktonic and biofilm cells of SBP strain . In the LBP, *FimH* increased in 7.6-fold and 8.3-fold in planktonic cells (Figure 5). Thus, *FimH* expression was not correlated with biofilm development of any strains. This observation is highly aligning with previous reports that demonstrate the function of *FimH* in colonization, biofilm formation and persistence in UTI, when compared to type 3 fimbriae, *FimH* plays least role in biofilm formation on abiotic as well as biotic surfaces (Di Martino et al. 2003a).

#### mrkD gene

Type 3 fimbriae (*mrkD*) gene expression was downregulated in biofilm cells of both strains (SBP and LBP). Hence playing no part in the formation of *Klebsiella* biofilms.

#### WabG and WcaG gene

*WabG* plays a vital role in the synthesis of core lipopolysaccharide and capsular polysaccharide of *K. pneumoniae.* In SBP and LBP strains, relatable to other genes, *WabG* were upregulated ten times higher both in planktonic and biofilm cells (Figure 5). When tested with *WcaG*, the expression was downregulated in both strains in planktonic and biofilm. Though the CT values of qPCR was observed in certain conditions but desired amplicon were not obtained in agarose gel.

In sum-up, we found the cations such as K^+^ and Ca^2+^ plays vital role on molecular level enhancing the biofilm formation through LPS by means of *WbaG* gene expression. Our observation in respect to capsule size and its related genes (*rmpA* and *WcaG*) expression with the biofilm development was independent. Similar, the fimbriae genes (*FimH* and *mrkD)* exhibit co-regulated expression.

## Discussion

Globally, infections caused by *K*. pneumoniae tend to be resistant to third generation carbapenem antibiotics (Temkin et al. 2018). Comparison to respiratory and blood, the urine strains are highly antibiotic resistant and leading to high morbidity and mortality world-wide (Ballén et al. 2021, Ikuta et al. 2022). Human urine is a rich bodily fluid with more than 3000 metabolites (carbon & nitrogen source) being an excellent seed, supporting the bacteria growth. At the same time, it is highly concentrated with antibacterial factor such as IgA (slgA), lactoferrin, D-serine, Urea and oligosaccharides. However, the bacteria have decoded environment very well over the years by developing a survivability factors. Such as siderophore to scavenger iron from lactoferrin, D-serine deaminase to degrade the serine residues as well as glycine betaine to confer resistance to high osmolarity occurred due to urea deposition. In this instance, the untreated UTI can progress to bacteraemia causing uroepithelial cell damage (Flores-Mireles et al. 2015). Increasing the mortality rate from 13.53% (UTI) to 54.30% in blood stream infections (Xu, Sun, and Ma et al. 2017). Among the various *Klebsiella* virulence factors, biofilm a dynamic shield causing over 80% chronic infection and more than 60% of bacterial infections (Moscoso, García, and López et al. 2009).

Conversely, mutation of these fimbriae (*FimH* and *MrkD*), LPS (*WabG*) and capsule (*WcaG* & *RmpA*) genes diminish *Klebsiella* biofilm. (Izquierdo et al. 2003, Schroll et al. 2010, Li et al. 2019, Zheng et al. 2018). Our findings indicated, in presence of glucose the ions such as Mg^2+,^ K^+^, Ca^2+,^ Na^+^, NH^+^_4_, and AS positively stimulated the biofilm development of all the strains. In the kinetic study, the biofilm was higher during 72 h in all conditions. When compared to clinical isolates, the environmental strain (kp4) produced a modest amount of biofilm in all examined medium. Among tested salts, K^+^ and Ca^2+^ enhanced higher biofilm (>65%) followed by Na^+^ (>50%) in the presences of ions. Increasing the concertation of Ca^2+^ ions affected the biofilm formation in all strains. Furthermore, we have discovered that the mass of the biofilm and the CFU of planktonic cells or capsule size are unrelated. Even, our investigation on colony morphology was not correlated with *Klebsiella* biofilm. However, A new observation was noticed, in the presence of monovalent ions (K^+^, NH^+^_4_, and Na^+^), pigmented colony morphology was observed in all the strains, whereas such colony morphology was absent in the presence of divalent ions (Ca^2+^ & Mg^2+^) (Suppl.Table 3).

Numerous studies on the promotion of biofilm by monovalent (K^+^ & Na^+^) and divalent (Ca^2+^) ions have been published for many bacteria species except in *Klebsiella*. Justified mechanism involves, K^+^ ions enhance the biofilm by means of cell communication (Jing et al. 2022). Na^+^ and Ca^2+^ ions bind with phosphate and carboxylate group of LPS regulating cell-cell and cell surface adhesion (Simoni et al. 2000). Furthermore, Ca^2+^ ion has been shown via extensive study to have multi-role in biofilms formation. i) *Salmonella enterica*, alter the adhesion protein SiiE increasing the adherence (Griessl et al. 2013), ii) Strengthen the adhesion protein by extending contact to the substratum, iii) Charge neutralization, bridging the cell & substratum and surface hydrophobicity aiding the bacterial attachment and increased thickness of *Pseudomonas mendocina* biofilm (He et al. 2016, Song and Leff 2006, Mangwani et al. 2014).

Nevertheless, the are no report on the role of ions over the biofilm related genes of *Klebsiella*. In alignment with previous report, we have also noticed a strict co-regulation system of *FimH* and *MrkD* in *K. pneumoniae* (Stahlhut et al. 2012). In contrast, we find no association between the production of biofilms and the expression of the fimbriae genes, type I and III, in biofilm cells. Apart from *Wcab* expression in glucose, no other capsule-related genes (*Wcab* and *RmpA*) were expressed in presence of cations. Whereas, the LPS component gene *WbaG* was highly expressed in all cation media, with the maximum fold of 16.198-fold in the presence of Ca^2+^, in contrast to the capsule and fimbriae genes. A direct correlation of increment in fold expression of *WbaG* upregulation and biofilm mass was clearly displayed in all the conditions. Stating, presence of ions in both urine as well as in blood stream, cause the production of LPS increasing the *Klebsiella* biofilm. We advocate that cations enhance the expression of LPS gene (*WbaG*), facilitating a potential *Klebsiella* biofilm by increasing the colonization and cell to cell interactions.

## Conclusion

The present study elucidated the role of cations on biofilm development of *Klebsiella* spp. The research demonstrated that Ca^2+^ and K^+^ ions, in contrast to other cations, stimulated the formation of biofilms in *Klebsiella* spp. Additionally, we have observed that *Klebsiella* sp. biofilm development is inhibited by increased Ca^2+^ concentrations, proving its role in molecular regulation rather than electrostatic interaction between the cell and surface. In the presence of all tested cations, there was no correlation between the cell density or capsule thickness and *Klebsiella* biofilm. Apart from capsule nor fimbriae rather the LPS synthesis gene (*WbaG*), plays a significant role in the formation of *Klebsiella* biofilm, suggesting as potent target in the treatment of *Klebsiella* biofilm in UTI.

## Acknowledgment

The authors express their gratitude to SASTRA Deemed University for providing financial assistance through the T.R.R grant (Ref. No: SASTRA-TRR-SCBT). and appreciative of the facilities that allowed us to conduct our research.

## Author’s contribution

AM performed all the experiments and analysed the results. SG performed biofilm development at different pH and Ca^2+^ concertation. JR obtained funding and designed the work. Manuscript was written and verified by AM & JR.

