## Supplemental table 3,4,5 for "Elucidating the effect of cations on *Klebsiella pneumoniae* biofilm and related genes"

**Supplementary**

**Table 3:** Morphology characteristics of the seven *K. pneumoniae* in presence Glucose, Mg^2+^, K^+^, NH^+^_4_, Na^+^, Ca^2+^, and AS cultured in congo red agar.

| **Strain** | **Cation ion** | **Form** | **Elevation** | **Margin** | **Pigment** |
| --- | --- | --- | --- | --- | --- |
| **Kp1** | Mg^2+^ | Irregular | Raised | Undulate | No |
|  | K^+^ | Irregular | Flat | Undulate | Black rim |
|  | NH^4+^ | Irregular | Umbo | Undulate | Red rim |
|  | Na^+^ | Irregular | Flat | Entire | Pink rim |
|  | Ca^2+^ | Irregular | Raised | Undulate | No |
|  | AS | Irregular | Flat | Undulate | Black rim |
| **Kp2** | Mg^2+^ | Circular | Raised | Entire | No |
|  | K^+^ | Circular | Umbo | Entire | Yes red rim |
|  | NH^4+^ | Circular | Convex | Entire | No |
|  | Na^+^ | Irregular | Flat | Undulate | No |
|  | Ca^2+^ | Circular | Raised | Entire | No |
|  | AS | Irregular | Flat | Undulate | No |
| **Kp3** | Mg^2+^ | Circular | Convex | Entire | No |
|  | K^+^ | Circular | Convex | Entire | No |
|  | NH^4+^ | Irregular | Umbo | Entire | Pink rim |
|  | Na^+^ | Irregular | Umbo | Undulate | Pink dots |
|  | Ca^2+^ | Irregular | Convex | Undulate | No |
|  | AS | Irregular | Raised | Undulate | Black rim |
| **Kp4** | Mg^2+^ | Circular | Flat | Entire | No |
|  | K^+^ | Nil | Nil | Nil | No |
|  | NH^4+^ | Irregular | Umbo | Undulate | Red rim |
|  | Na^+^ | Nil | Nil | Nil | No |
|  | Ca^2+^ | Irregular | Flat | Undulate | No |
|  | AS | Circular | Raised | Entire | Pink in entire |
| **Kp5** | Mg^2+^ | Circular | Convex | Entire | No |
|  | K^+^ | Circular | Raised | Entire | No |
|  | NH^4+^ | Irregular | Umbo | Undulate | Pink rim but slight |
|  | Na^+^ | Irregular | Umbo | Undulate | No |
|  | Ca^2+^ | Irregular | Raised | Undulate | No |
|  | AS | Irregular | Raised | Entire | No |
| **BA 33560** | Mg^2+^ | Circular | Convex | Entire | No |
|  | K^+^ | Irregular | Raised | Undulate | No |
|  | NH^4+^ | Nil | Nil | Nil | No |
|  | Na^+^ | Irregular | Flat | Entire | Pink dots |
|  | Ca^2+^ | Irregular | Raised | Entire | No |
|  | AS | Circular | Raised | Entire | Black rim |
| **BA 28434** | Mg^2+^ | Irregular | Raised | Undulate | No |
|  | K^+^ | Circular | Flat | Entire | Black rim |
|  | NH^4+^ | Circular | Flat | Entire | Red rim |
|  | Na^+^ | Punctiform | Umbo | Undulate | Pink dots |
|  | Ca^2+^ | Irregular | Flat | Undulate | No |
|  | AS | Circular | Umbo | Entire | Black rim |

**Table 4**: Artificial urine media composition

| Component | Quantity (g) | Concentration (mMol l^-1^) |
| --- | --- | --- |
| Peptone L37 | 1 |  |
| Yeast extract | .005 |  |
| Lactic acid | 0.1 | 1.1 |
| Citric acid | 0.4 | 2 |
| Sodium bicarbonate | 2.1 | 25 |
| Urea | 10 | 170 |
| Uric acid | 0.07 | 0.4 |
| Creatinine | 0.8 | 7 |
| Calcium choride.2H2O | 0.37 | 2.5 |
| Sodium chloride | 5.2 | 90 |
| Iron II sulphate.10H2O | 0.0012 | 0.005 |
| Magnesium sulphate.7H_2_O | 0.49 | 2 |
| Sodium sulphate.7H_2_O | 3.2 | 10 |
| Potassium dihydrogen phosphate | 0.49 | 7 |
| di-potassium hydrogen phosphate | 1.2 | 7 |
| Ammonium chloride | 1.3 | 25 |
| Distilled water | 11 |  |

**Table 5**: Cell size, capsule size and Capsule percentage of the seven *K. pneumoniae* in presence Glucose, Mg^2+^, K^+^, NH^+^_4_, Na^+^, Ca^2+^, and AS.

| **Strain** | **Cation ion** | **Cell size (mm)** | **Capsule (mm)** | **Capsule%** |
| --- | --- | --- | --- | --- |
| **Kp1** | G | 8.141 | 2.449 | 23.12559 |
|  | Mg^2+^G | 3.388 | 3.542 | 51.11111 |
|  | K^+^+G | 9.168 | 3.9338 | 30.02488 |
|  | NH_4_+G | 7.601 | 1.657 | 17.89803 |
|  | Na^+^+G | 10.665 | 0.747 | 6.545741 |
|  | Ca^2+^+G | 6.838 | 2.335 | 25.45514 |
|  | AS+G | Nil | Nil | Nil |
| **Kp2** | G | 8.876 | 1.424 | 13.82524 |
|  | Mg^2+^G | 6.935 | 1.903 | 21.53202 |
|  | K^+^+G | 8.746 | 0.828 | 8.648423 |
|  | NH_4_+G | 8.247 | 0.561 | 6.36921 |
|  | Na^+^+G | 9.387 | 0.006 | 0.063877 |
|  | Ca^2+^+G | 3.252 | 2.734 | 45.67324 |
|  | AS+G | 4.376 | 2.074 | 32.15504 |
| **kp3** | G | 7.964 | 2.102 | 20.88218 |
|  | Mg^2+^G | 9.66 | 1.346 | 12.22969 |
|  | K^+^+G | 7.801 | 1.45 | 15.67398 |
|  | NH_4_+G | Nil | Nil | Nil |
|  | Na^+^+G | 5.81 | 1.536 | 20.90934 |
|  | Ca^2+^+G | 6.16 | 1.157 | 15.81249 |
|  | AS+G | 9.752 | 0.393 | 3.873829 |
| **Kp4** | G | 8.213 | 1.055 | 11.38325 |
|  | Mg^2+^G | 6.486 | 0.331 | 4.855508 |
|  | K^+^+G | Nil | Nil | Nil |
|  | NH_4_+G | 8.793 | 2.372 | 21.24496 |
|  | Na^+^+G | 13.003 | 2.925 | 18.36389 |
|  | Ca^2+^+G | 6.639 | 1.986 | 23.02609 |
|  | AS+G | 7.883 | 2.038 | 20.54228 |
| **Kp5** | G | Nil | Nil | Nil |
|  | Mg^2+^G | 4.333 | 0.338 | 7.236138 |
|  | K^+^+G | 5.757 | 0.544 | 8.63355 |
|  | NH_4_+G | 4.807 | 0.309 | 6.039875 |
|  | Na^+^+G | 3.925 | 0.449 | 10.2652 |
|  | Ca^2+^+G | 4.724 | 0.84 | 15.09705 |
|  | AS+G | 8.37 | 1.834 | 17.97334 |
| **BA 33560** | G | 8.735 | 1.242 | 12.44863 |
|  | Mg^2+^G | 7.869 | 2.431 | 23.60194 |
|  | K^+^+G | 4.845 | 0.712 | 12.81267 |
|  | NH_4_+G | 8.172 | 0.32 | 3.768252 |
|  | Na^+^+G | 4.117 | 0.914 | 18.16736 |
|  | Ca^2+^+G | 9.428 | 1.932 | 17.00704 |
|  | AS+G | 12.389 | 1.292 | 9.443754 |
| **BA 28434** | G | 9.415 | 1.43 | 13.1858 |
|  | Mg^2+^G | 6.774 | 1.138 | 14.38322 |
|  | K^+^+G | 7.7 | 2.197 | 22.19865 |
|  | NH_4_+G | 8.558 | 0.427 | 4.752365 |
|  | Na^+^+G | 8.336 | 3.576 | 30.02015 |
|  | Ca^2+^+G | 8.571 | 1.863 | 17.85509 |
|  | AS+G | 9.198 | 1.929 | 17.33621 |
